# Rapamycin-sensitive mechanisms confine the growth of fission yeast below the temperatures detrimental to cell physiology

**DOI:** 10.1101/2023.05.04.539340

**Authors:** Yuichi Morozumi, Fontip Mahayot, Yukiko Nakase, Jia Xin Soong, Sayaka Yamawaki, Fajar Sofyantoro, Yuki Imabata, Arisa H. Oda, Miki Tamura, Shunsuke Kofuji, Yutaka Akikusa, Kunihiro Ohta, Kazuhiro Shiozaki

**Affiliations:** Division of Biological Science, Nara Institute of Science and Technology, Ikoma, Nara 630-0192, Japan; Department of Animal Physiology, Faculty of Biology, Universitas Gadjah Mada, Yogyakarta 55281, Indonesia; Department of Life Sciences, Graduate School of Arts and Sciences, The University of Tokyo, Meguro-ku, Japan; Department of Microbiology and Molecular Genetics, University of California, Davis, CA 95616, USA

## Abstract

Cells cease to proliferate above their growth-permissible temperatures, a ubiquitous phenomenon generally attributed to protein denaturing and heat damage to other cellular macromolecules. We here report that, in the presence of the macrolide compound rapamycin, the fission yeast *Schizosaccharomyces pombe* can proliferate at high temperatures that normally arrest its growth. Rapamycin is a potent inhibitor of the protein kinase complex TOR Complex 1 (TORC1), and consistently, mutations to the TORC1 subunit RAPTOR/Mip1 and the TORC1 substrate Sck1 significantly improve cellular heat resistance. These results suggest that TORC1, a well-established growth promoter, restricts the high-temperature growth of fission yeast and that compromised TORC1 signaling allows cell proliferation at higher temperatures. Aiming for a more comprehensive understanding of the negative regulation of high-temperature growth, we conducted genome-wide screens in *S. pombe*, which identified Sck1 and additional factors that appear to suppress cell proliferation at high temperatures. Our study has uncovered unexpected mechanisms of growth restraint even below the temperatures deleterious to cell physiology. Thus, growth arrest at high temperatures may not directly result from heat damage to cellular components essential for proliferation and viability.

**Significance Statement:** The immunosuppressant rapamycin is a specific inhibitor of the protein kinase Target Of Rapamycin (TOR), and the drug is known to extend the lifespan of diverse eukaryotic organisms. In this study, we have found that rapamycin confers heat resistance on fission yeast, allowing its proliferation above the normal permissive temperatures. This unexpected observation suggests that TOR, which is known as a growth-promoting kinase, is inhibitory to cell proliferation at high temperatures. Our genome-wide screens have identified additional genes whose deletion leads to improved growth under heat stress. Thus, cells may have mechanisms that restrict proliferation even below the temperatures deleterious to their physiology.

## Introduction

Although the habitat temperatures of living organisms are surprisingly diverse, each species has a relatively narrow range of optimal growth temperatures. Even a moderate rise in the growth temperatures can cause heat stress detrimental to the stability and/or activity of proteins (1, 2). As counteracting measures, cells activate the heat shock response, an evolutionarily conserved program that induces expression of the molecular chaperons termed heat shock proteins (HSPs) (3, 4). The molecular chaperones maintain cellular proteostasis through refolding of denatured proteins, prevention of their aggregation, and sorting of unfolded proteins that are marked for degradation (3, 5). As the accumulation of misfolded protein aggregates is potentially harmful to cells (2, 6), molecular chaperones have been extensively studied as a major pro-survival mechanism in response to acute heat shock. On the other hand, it remains unclear whether proteotoxic stress is responsible for compromised or arrested growth of cells only marginally above their survivable temperatures.

The Target of Rapamycin (TOR), a member of the phosphoinositide-3 kinase-related kinase (PIKK) family, coordinates eukaryotic cell growth and proliferation with environmental cues, including nutrients and growth factors (7). TOR functions by forming two distinct complexes, namely TOR complex 1 (TORC1) and TOR complex 2 (TORC2). The immunosuppressant rapamycin forms an intracellular complex with the FKBP12 protein, which then binds to the TOR kinase in TORC1 but not in TORC2, thereby leading to the specific inhibition of TORC1 (8, 9). In response to nutrient and growth stimuli, TORC1 modulates anabolic processes, including the synthesis of proteins, nucleotides and lipids, as well as cellular catabolism such as autophagy (10, 11).

Mammalian TORC1 (mTORC1), whose subunits include RAPTOR and mLST8, is regulated by two classes of small GTPases, RHEB and RAGs (12, 13). Upon amino-acid stimuli, the active form of the RAG heterodimer (GTP-bound RAGA/B with GDP-bound RAGC/D) recruits mTORC1 to lysosomal membranes, where GTP-bound, active RHEB activates mTORC1 through direct interaction. (14, 15). Activated mTORC1 then phosphorylates multiple substrates to promote cell growth. Ribosomal protein S6 kinase 1 (S6K1), an AGC-family kinase, and eukaryotic initiation factor 4E-binding protein 1 (4EBP1) are the best-characterized mTORC1 substrates that regulate mRNA translation (16–19).

The TORC1 signaling pathway is highly conserved between mammalian cells and the fission yeast *Schizosaccharomyces pombe* (20). In *S. pombe*, the catalytic subunit Tor2 forms TORC1 together with the RAPTOR equivalent Mip1 and Wat1/Pop3, a mLST8 ortholog (21–23). The fission yeast RHEB ortholog termed Rhb1 functions as an essential activator of TORC1 (24–26), while TORC1 is negatively regulated by the Gtr1-Gtr2 heterodimer, which is a counterpart of the mammalian RAGA/B-RAGC/D complex (27, 28). Psk1, an AGC-family kinase orthologous to mammalian S6K1, has been identified as a substrate of TORC1 (29, 30). TORC1 is essential for cell growth, and the loss of the TORC1 function leads to cell cycle arrest in G1 both in budding and fission yeasts (8, 21–23, 26, 31). On the other hand, suppression of TORC1 by rapamycin brings about no apparent growth inhibition in fission yeast (32), in stark contrast to the observation in budding yeast (33); rapamycin appears to only partially inhibit the TORC1 activity in *S. pombe* (34).

The optimal growth temperature of *S. pombe* cells is around 30°C; they exhibit somewhat impaired growth at 37°C and fail to grow at 38°C and above. We have found that fission yeast cells remain viable and proliferate even at 39°C when the TORC1 activity is suppressed by rapamycin. Moreover, the null mutant of the Sck1 kinase, a known TORC1 substrate (29, 35), exhibits significant cell growth at 39°C even in the absence of rapamycin. Thus, unexpectedly, the growth of fission yeast cells at high temperatures appears to be negatively regulated by TORC1, which has been established as a growth promoter in *S. pombe* and other eukaryotes. Furthermore, our genome-wide screens have identified several additional factors that appear to restrict cellular growth at high temperatures. These findings strongly suggest that the high-temperature growth of fission yeast is impeded by inherent mechanisms even below the temperatures detrimental to cell physiology. Although the heat sensitivity of cell growth is a ubiquitous phenomenon, it may not necessarily be due to proteotoxic stress or other heat-induced damage to cellular components.

## Results

### Suppression of TORC1 activity promotes *S. pombe* growth at high temperatures

In the fission yeast *S. pombe*, the optimal growth temperature of wild-type cells is around 30°C, and their growth is significantly impaired at 38°C and completely ceased at 39°C or higher (Fig. 1A, left). We serendipitously discovered that fission yeast cells exhibit significant proliferation even at 38°C and 39°C, but not at 40°C, on agar medium supplemented with rapamycin, a TORC1-specific inhibitor (Fig. 1A, right). Such enhanced growth at 39°C was also observed when culturing *S. pombe* cells in liquid medium with rapamycin, where inhibition of the TORC1-dependent phosphorylation of the S6K1 ortholog Psk1 (29) was confirmed (Fig. S1). The cell viability assay with methylene blue staining (36, 37) indicated that, at 39°C, cell viability was largely lost by 48 hours, which was counteracted by rapamycin (Fig. 1B).

**Fig. 1.**
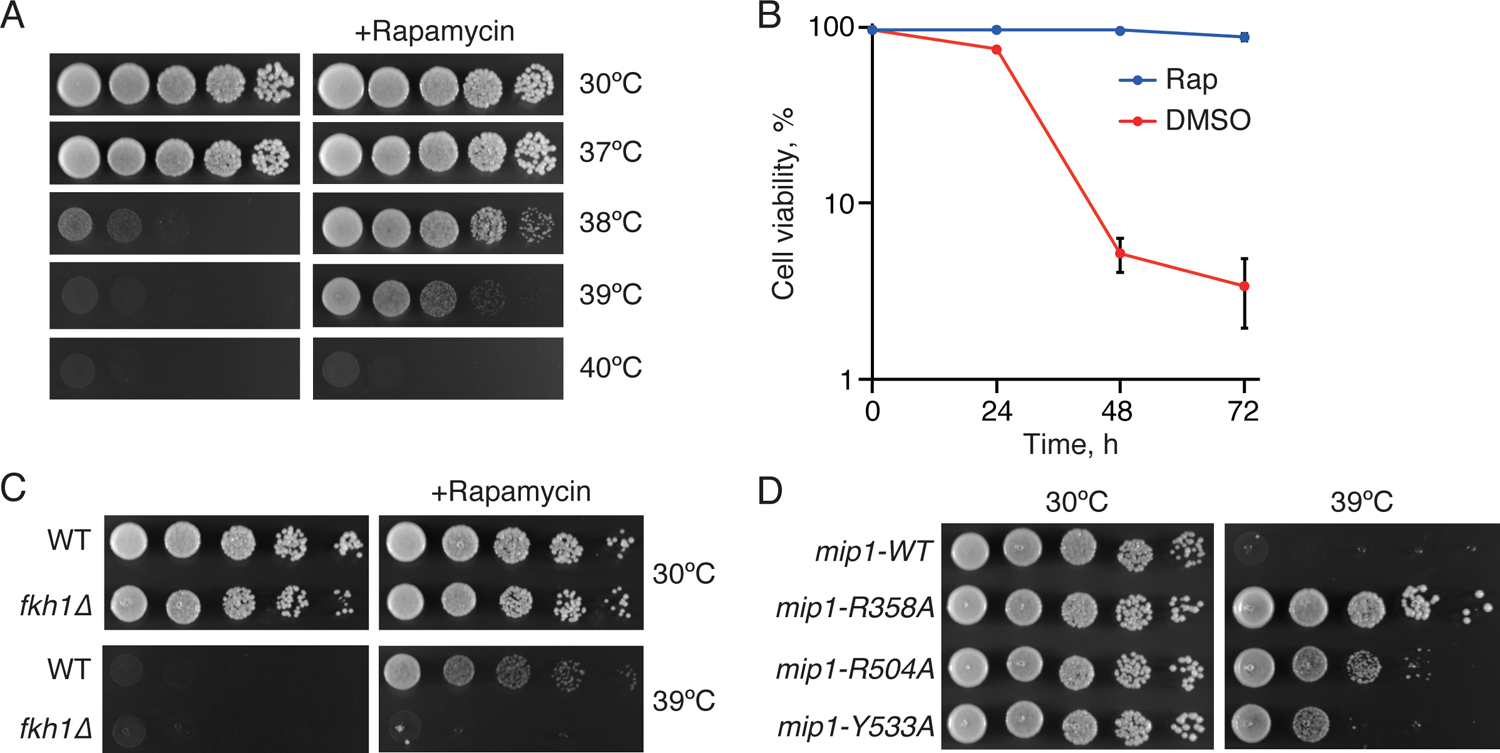
Suppression of TORC1 activity allows *S. pombe* cells to grow at high temperatures. (A) Wild-type cells were grown in YES liquid medium at 30°C. Their serial dilutions were spotted onto YES agar plates in the presence and absence of 100 ng/ml rapamycin for growth assay at indicated temperatures. (B) Wild-type cells were grown in YES liquid medium at 30°C. At time 0, the culture concentration was adjusted to OD_600_ = 0.1 ± 0.05, and the cell viability in the presence (Rap) and absence (DMSO) of 200 ng/ml rapamycin at 39°C was measured by methylene blue staining at indicated time points. Data are presented as means ± s.d. from three independent experiments, and at least 200 cells were counted for each experiment. (C) The *fkh1Δ* mutant failed to grow at 39°C even in the presence of rapamycin. The growth of wild-type (WT) and *fkh1Δ* cells was tested as in (A). (D) The *mip1* mutants exhibit cell proliferation at 39°C even in the absence of rapamycin. The indicated *mip1* mutant strains were grown in YES medium at 30°C, and their serial dilutions were spotted onto YES agar plates for growth assay at 30°C and 39°C.

The data described above suggest that suppression of TORC1 activity by rapamycin confers heat resistance on fission yeast cells. To further corroborate the possibility, the growth of a mutant strain lacking Fkh1, a FKBP12 ortholog that mediates rapamycin-dependent inhibition of TORC1 (38–40), was examined at high temperatures. As expected, the *fkh1* null mutant failed to grow at 39°C even in the presence of rapamycin (Fig. 1C), consistent with the idea that the rapamycin-induced heat resistance is due to TORC1 inhibition. Furthermore, even in the absence of rapamycin, mutations to the TORC1 subunit Mip1, a RAPTOR ortholog (21–23), allowed fission yeast cells to grow at high temperatures (Fig. 1D). Strains carrying *mip1-R358A*, *mip1-R504A*, or *mip1-Y533A* mutations have attenuated TORC1 activity (30), and they grew at 39°C where *mip1*^+^ cells failed to do so.

Taken together, these results indicate that suppression of TORC1 activity allows *S. pombe* cells to remain viable and proliferate at growth-inhibitory high temperatures up to 39°C.

### TORC1-Sck1 signaling suppresses cellular growth at high temperatures

The results above imply that TORC1 activity somehow restricts cell growth at high temperatures. To further explore the events downstream of TORC1 in the control of high-temperature growth, we examined the phenotypes of the null mutants that lack the known TORC1 substrates in fission yeast; the AGC-family kinases Psk1, Sck1 and Sck2 (29), the RNA polymerase III repressor Maf1 (41), and the autophagy regulator Atg13 (35). Among them, only the *sck1* null mutant (*sck1Δ*) exhibited growth at 39°C even in the absence of rapamycin (Fig. 2A). A similar heat-resistant phenotype was observed with a strain expressing the catalytically inactive Sck1 kinase, Sck1KD, in which Lys-331 in the ATP-binding domain is substituted with alanine (Fig. 2B). These results suggest that TORC1-dependent activation of the Sck1 kinase negatively regulates cell growth at high temperatures.

**Fig. 2.**
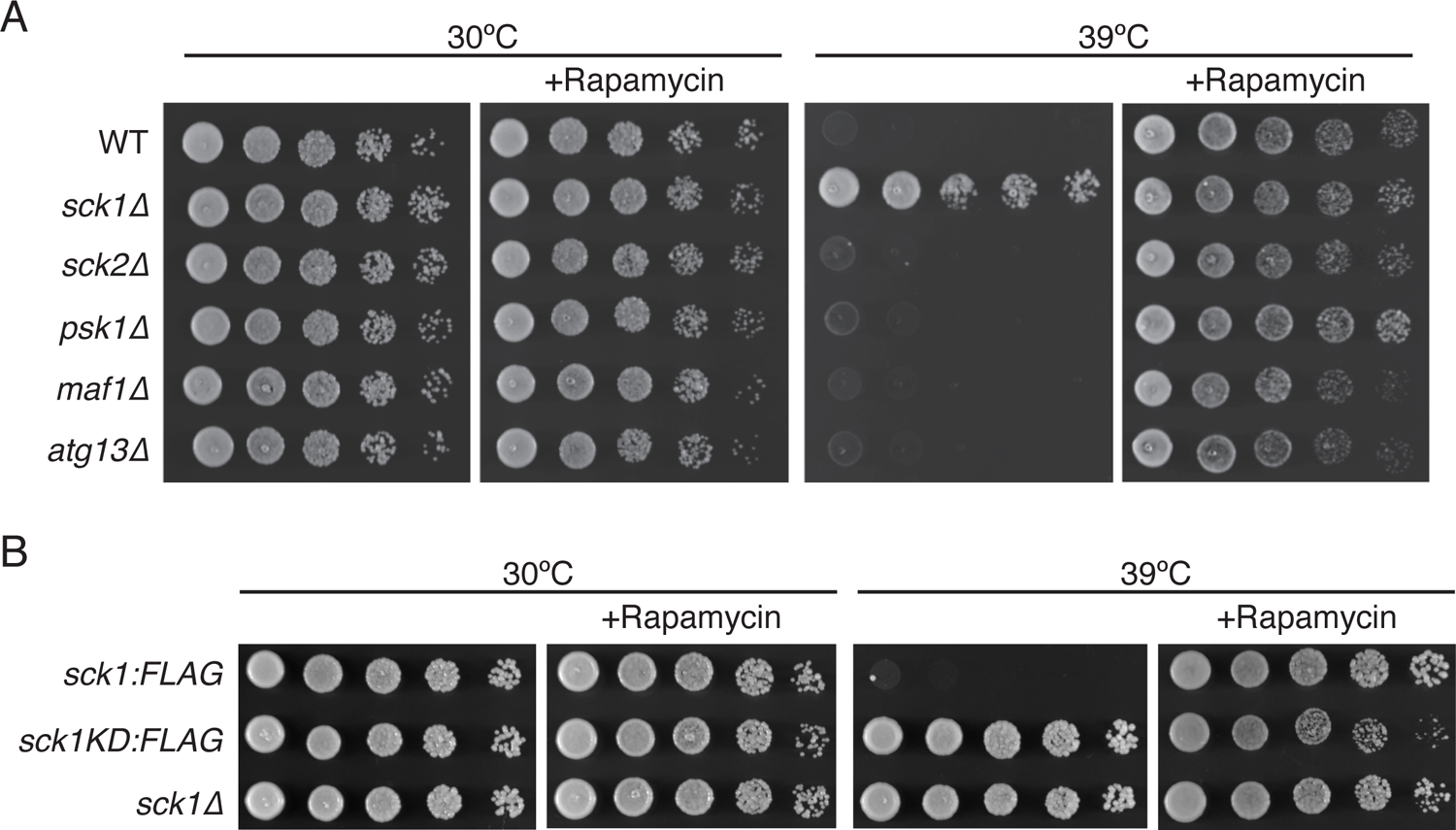
The Sck1 kinase inhibits cell growth at high temperatures. (A) Wild-type (WT) and the indicated null mutant cells were grown in YES liquid medium at 30°C. Their growth in the presence and absence of rapamycin (100 ng/ml) was tested at 30 and 39°C by spotting the serial dilutions of their cultures on YES agar plates. (B) The catalytic activity of Sck1 is required for suppressing high-temperature growth. The growth of the *sck1:FLAG*, *sck1-K311A:FLAG* and *sck1Δ* strains was tested as in (A).

### A genome-wide screen for genes inhibitory to cellular growth at high temperatures

Our findings described above revealed that *S. pombe* has mechanisms to proactively suppress cell growth at high temperatures, such as 39°C, including TORC1-Sck1 signaling. For more comprehensive identification of the regulatory elements that restrict high-temperature growth, we screened a haploid null mutant library of approximately 3,400 nonessential genes for mutants that can grow at 39°C. Not surprisingly, the screen re-isolated the *sck1Δ* mutant, along with the null mutants of SPAC1420.01c, *dri1*^+^, SPBC17D1.05, and *mtq2*^+^ (Fig. 3A). Open reading frame SPAC1420.01c encodes a protein orthologous to budding yeast Mks1 and therefore, this fission yeast protein is referred to as Mks1 hereafter. The Mks1 orthologs in these two yeast species share limited overall sequence homology with a conserved short stretch in the N-terminal region (Fig. S2). The growth of the *S. pombe mks1*Δ mutant at 39°C was comparable to that of *sck1*Δ and better than that of the *dri1*Δ, whose growth phenotype was similar to that of *SPBC17D1.05Δ* (Fig. 3A). Dri1 is a putative RNA-binding protein involved in heterochromatin assembly and kinesin loading to the mitotic spindle (42, 43), while SPBC17D1.05 is an uncharacterized, fission yeast-specific protein, which we named Rhs1 (<u>R</u>egulator of <u>h</u>eat <u>s</u>ensitivity). Among the isolated mutants, only *mtq2Δ* cells showed compromised growth even at 30°C, though they also grew at 39°C (Fig. 3A). Mtq2 is predicted to be the catalytic subunit of the eRF1 methyltransferase complex; its ortholog in *S. cerevisiae* forms a heterodimer with Trm112 to methylate the translation release factor eRF1, which is essential for translation termination (44, 45).

**Fig. 3.**
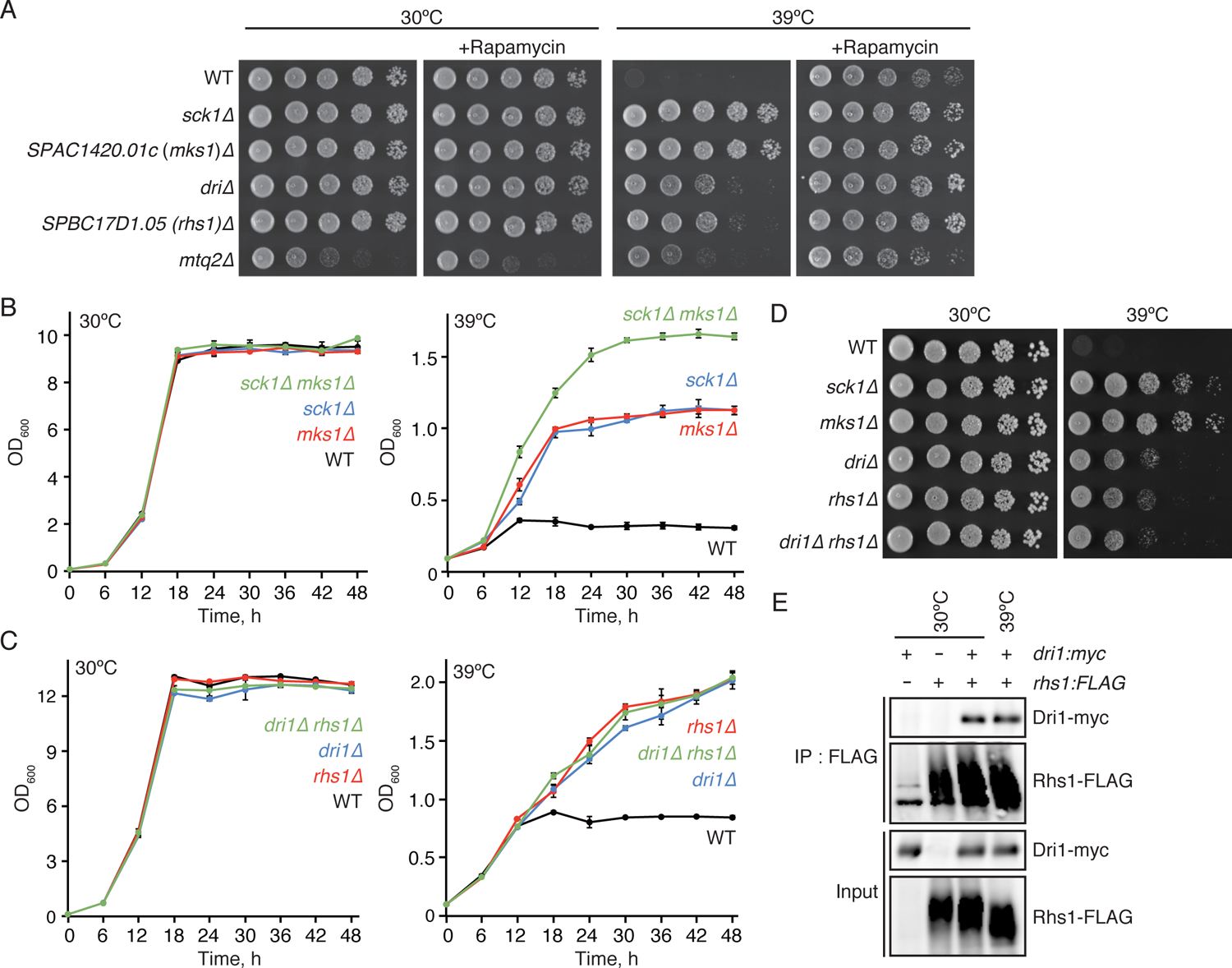
Identification of genes that suppress cellular growth at high temperatures. (A) Isolated gene deletion mutants that exhibit growth at high temperatures. The indicated null mutants were grown in YES medium at 30°C. Their growth in the presence and absence of rapamycin (100 ng/ml) was tested at 30 and 39°C by spotting serial dilutions of their cultures on YES agar plates. (B) Sck1 and Mks1 function independently to inhibit cell growth at high temperatures. Wild-type (WT) and the indicated mutant cells were cultured in YES medium at 30°C. At time 0, the culture concentration was adjusted to OD_600_ = 0.1 ± 0.05, and their growth at 30 and 39 was monitored by measuring OD_600_ every 6 hours. Data are presented as means ± s.d. from three independent experiments. (C, D) Dri1 and Rhs1 function in the same pathway. (C) The growth kinetics of indicated mutant strains were measured as in (B). (D) The indicated mutant strains were grown in YES medium at 30°C, and their serial dilutions were spotted onto YES agar plates for growth assay at 30°C and 39°C. (E) Dri1 physically interacts with Rhs1. Strains expressing Rhs1-FLAG and Nrp1-*myc* individually or simultaneously were grown in YES medium at 30°C and 39°C, and the cell lysate was subjected to immunoprecipitation using anti-FLAG affinity beads (IP: FLAG). The samples were analyzed by anti-FLAG and anti-*myc* immunoblotting.

In order to examine the functional relationships among the genes identified in the screen, their double mutants were constructed for epistasis analyses. Growth kinetics monitored in liquid medium at 30°C found no apparent difference among the wild-type, *mks1Δ*, *sck1Δ*, and *sck1Δ mks1Δ* strains (Fig. 3B). At 39°C, the deletion of the two genes showed an additive effect, and the *sck1Δ mks1Δ* double mutant grew better than the individual single mutants, suggesting that Mks1 and Sck1 function independently to negatively regulate cell proliferation at high temperatures. Similar additive effects were also observed when the *sck1Δ* or *mks1Δ* mutations were combined with either *dri1Δ* or *rhs1Δ* (Fig. S3). On the other hand, the growth of the *dri1Δ rhs1Δ* double mutant at 39°C was comparable to that of the respective single mutants when cultured in liquid medium (Fig. 3C) as well as on solid agar medium (Fig. 3D). These observations suggest that Dri1 and Rhs1 function in the same pathway, but independently of Sck1 and Mks1. Interestingly, a fission yeast interactome study using the yeast two-hybrid assay detected the interaction between Dri1 and Rhs1 (46). To examine whether Dri1 forms a complex with Rhs1, we constructed a strain where the chromosomal *dri1*^+^ gene is tagged with the *myc* epitope sequence, and the Rhs1 protein is expressed with the FLAG epitope from the *rhs1^+^* locus. Immunoprecipitation of FLAG-tagged Rhs1 resulted in the co-purification of Dri1 at 30°C and 39°C (Fig. 3E). It is therefore likely that Dri1 and Rhs1 function as a complex in the suppression of cell growth at high temperatures.

### A mutation to the 14-3-3 protein Rad24, which forms a complex with Mks1, confers heat resistance on *S. pombe* cells

In addition to screening the gene deletion library as described above, we also attempted to isolate spontaneous mutations that confer heat resistance to fission yeast cells. When a wild-type strain was incubated at 39°C on agar growth medium, viable mutant colonies appeared at a frequency of ∼1×10^-5^, which were subsequently analyzed by whole genome sequencing. Though most heat-resistance mutations were found in the *sck1* or *dri1* genes, whose deletion also allows cell growth at 39°C (Fig. 3A), one mutant carried a single nucleotide substitution in *rad24*, a gene encoding a 14-3-3 protein. This mutation is predicted to replace Glu-185 with valine in Rad24, and the identical *rad24-E185V* mutation introduced to a wild-type strain by site-directed mutagenesis confirmed its ability to allow cell growth at 39°C (Fig. 4A). On the other hand, the *rad24* (*rad24Δ*) null mutant exhibited little growth at 39°C while growing better than wild-type cells at 38°C; thus, *rad24-E185V* does not appear to be a complete loss-of-function mutation. It should also be noted that in the presence of rapamycin, *rad24-E185V* and *rad24Δ* cells did not grow as much as wild-type cells at 39°C (Fig. 4A), implying a role of Rad24 in the rapamycin-induced heat resistance.

**Fig. 4.**
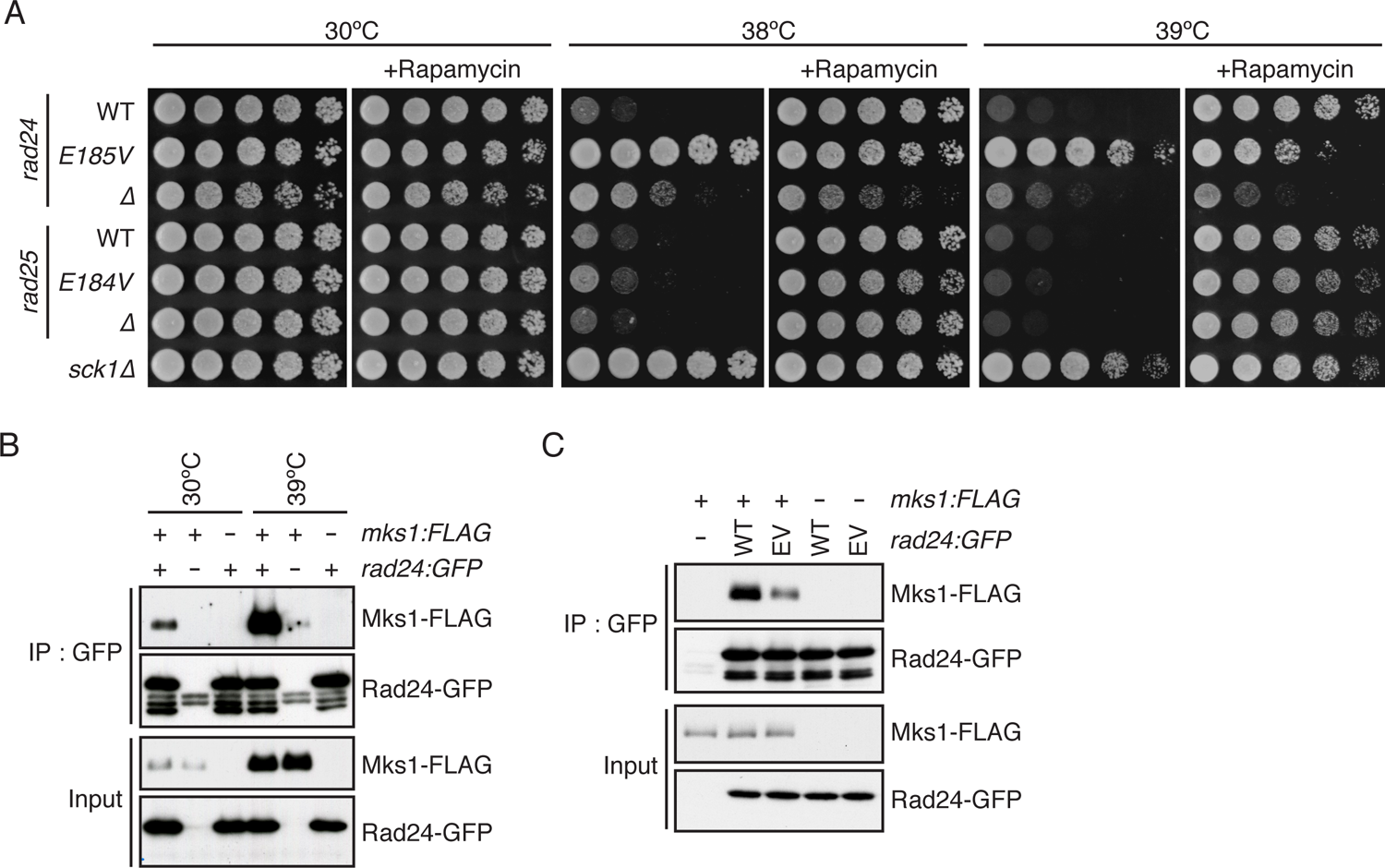
The *rad24-E185V* mutation, which compromises the Rad24-Mks1 interaction, confers heat resistance on *S. pombe* cells. (A) The *rad24-E185V* mutation confers heat resistance on fission yeast cells. The indicated strains were cultured at 30°C. Their growth in the presence and absence of rapamycin (100 ng/ml) was tested at indicated temperatures by spotting serial culture dilutions on YES agar plates. (B) Mks1 physically interacts with Rad24. Strains expressing Mks1-FLAG and Rad24-GFP individually or simultaneously were grown in YES medium at 30°C and sifted to 39°C. After 2-hour incubation at 39°C, the cell lysate was prepared, and Rad24-GFP was immunopurified using the antibodies against GFP (IP: GFP). Co-purified Mks1-FLAG was analyzed by immunoblotting. (C) The *rad24-E185V* mutation compromises the interaction of Rad24 with Mks1. *mks1-FLAG* cells expressing either Rad24 or Rad24-E185V tagged with GFP were grown, and their cell lysate was subjected to immunoprecipitation followed by immunoblotting as in (B).

Glu-185 in Rad24 is conserved in the *S. pombe* 14-3-3 protein paralog Rad25 as Glu-184 (Fig. S4). In contrast to the *rad24* mutants, both *rad25-E184V* and *rad25Δ* mutants failed to grow at 38°C and 39°C (Fig. 4A), suggesting little or no contribution of Rad25 to the growth regulation under high-temperature conditions.

Budding yeast Mks1 physically interacts with Bmh1, a 14-3-3 protein orthologous to fission yeast Rad24 (47). As our study in fission yeast identified Mks1 and Rad24 as negative regulators of high-temperature growth, we examined the physical interaction between Mks1 and Rad24. Immunoprecipitation of Rad24 tagged with the green fluorescent protein (GFP) resulted in co-purification of the FLAG-tagged Mks1 protein both at 30 and 39°C (Fig. 4B). A less amount of Mks1 was co-purified with Rad24-E185V than with the wild-type Rad24 protein (Fig. 4C). These results indicate that, like its ortholog in budding yeast, fission yeast Mks1 interacts with Rad24 and that their interaction is compromised by the *rad24-E185V* mutation.

### The Mks1 protein accumulates in cells exposed to high temperatures

In the experiments shown in Fig. 4B, we noticed that the amount of Mks1 in cells incubated at 39°C was significantly higher than that at 30°C, resulting in more Mks1 co-purified with Rad24 at 39°C. We confirmed that the Mks1 protein accumulated after the temperature shift from 30°C to 39°C and that rapamycin further promoted the accumulation (Fig. 5A); even at 30°C, rapamycin can induce the accumulation of Mks1 (Fig. S5A). On the other hand, the *mks1^+^* transcript did not significantly increase even after 4-hour incubation at 39°C (Fig. 5B), in contrast to the heat shock protein genes such as *hsp16^+^* (48). Thus, the accumulation of Mks1 at high temperatures does not appear to be due to the transcriptional regulation of *mks1*^+^. In addition, we observed a significantly elevated level of the Mks1 protein in the *mts2-1* background, a temperature-sensitive, hypomorphic allele of a 26S proteasome subunit (49), even at the permissive temperature (Fig. 5C). It is likely that proteasomal degradation regulates the cellular level of Mks1.

**Fig. 5.**
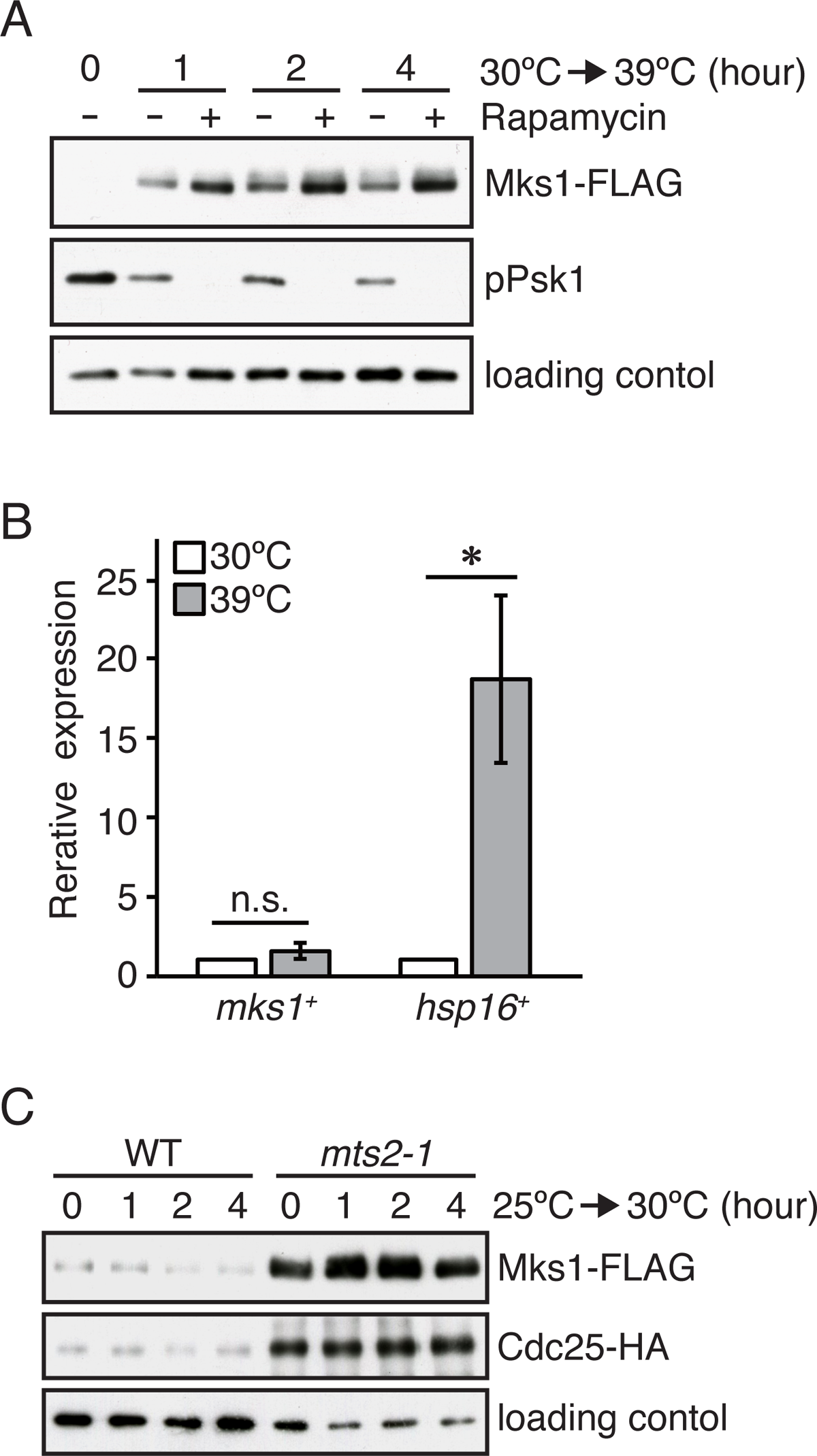
Cellular Mks1 protein accumulates upon heat stress by attenuation of its proteasomal degradation. (A) Heat stress induces the accumulation of Mks1, which is further enhanced by rapamycin. Cells were grown in YES medium at 30°C and shifted to 39°C in the presence and absence of 200 ng/ml rapamycin. Their cell lysate was probed with anti-FLAG and anti-Spc1 (loading control) antibodies. (B) Heat stress does not significantly change the transcription of the *mks1^+^* gene. The mRNA expression of *mks1^+^* and *hsp16^+^* was examined by qRT-PCR using cells grown in YES medium at 30°C and 39°C for 4 hours. The expression level of each gene is presented as a value relative to that at 30°C. Bars indicate ± s.d. from two independent experiments in triplicate. \**P*<0.05; n.s., not significant, compared to the mRNA expression at 30°C using Student’s t-test. (C) The protein accumulation of Mks1 in the *mts2-1* mutant. Wild-type (WT) and *mts2-1* mutant cells were grown in YES medium at 25°C and shifted to 30°C. Cells were harvested at indicated time points, and their cell lysate was subjected to immunoblotting using anti*-*FLAG, anti-HA, and anti-Spc1 (loading control) antibodies. Cdc25, known to accumulate in the *mts2-1* mutant (82), was utilized as a positive control.

It was previously observed in budding yeast that the 14-3-3 protein Bmh1 binds to Mks1 and protects it from proteasomal degradation (50). We therefore examined the cellular level of fission yeast Mks1 in the *rad24* mutants. As in wild-type cells, no accumulation of Mks1 was observed at 30°C in both *rad24Δ* and *rad24-E185V* mutants (Fig. S5B). In addition, the accumulation of Mks1 after the temperature shift to 39°C was not notably affected by the *rad24* mutations when compared to the wild-type. Thus, the 14-3-3 protein Rad24 does not have a major regulatory role in the proteasomal degradation of fission yeast Mks1.

### Mks1 has a conserved C-terminal motif required for the suppression of cell growth at high temperatures

As shown above, Mks1 interacts with the 14-3-3 protein Rad24, of which mutation, *rad24-E185V*, compromises their interaction and induces cellular heat resistance (Fig. 4). Therefore, we further delved into the Mks1-Rad24 interaction and its role in heat-resistant cell growth. To determine the Rad24-binding region within Mks1, we repeated the Rad24 immunoprecipitation assays with fission yeast strains expressing a series of truncated Mks1 proteins. Truncation of the N-terminal 300 residues of Mks1 (301-580) abrogated its interaction with Rad24, whereas Mks1 lacking the N-terminal 100 residues (101-580) or 200 residues (201-580) was found to associate with Rad24 (Fig. 6A). We also tested the mutant Mks1 proteins truncated from the C-terminus and found that Mks1(1-280) interacted with Rad24, implying that residues 201-280 of Mks1 are involved in the interaction with Rad24. As shown in Figure 6B, a mutant strain expressing Mks1(301-580) that cannot interact with Rad24 exhibited moderate cell growth at 39°C, whereas cells expressing the full length, 101-580, or 201-580 of Mks1 failed to grow at 39°C. These observations appear to be consistent with the idea that the Mks1-Rad24 interaction plays a role in the negative regulation of high-temperature growth.

**Fig. 6.**
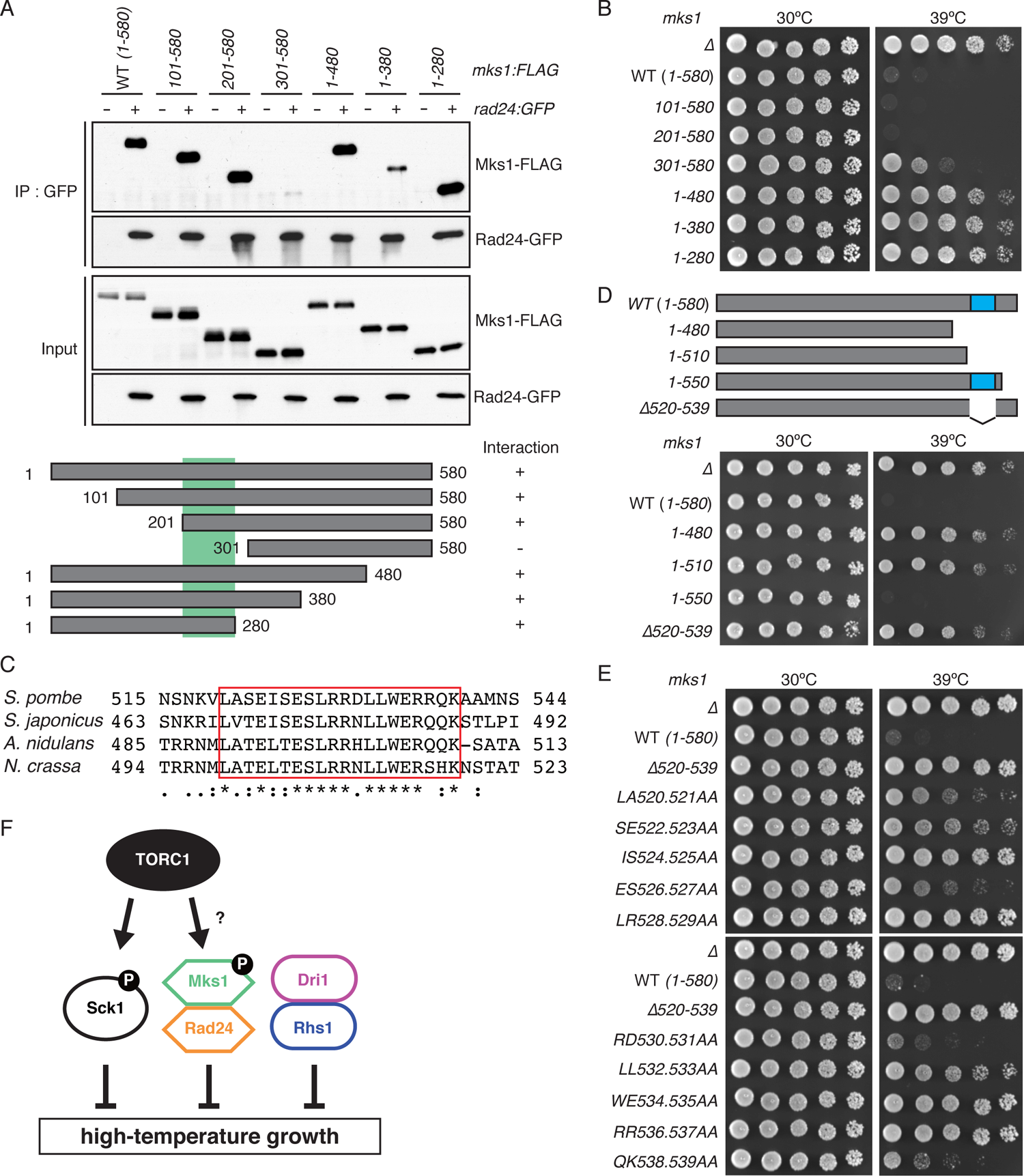
The C-terminal conserved region of Mks1 is indispensable for the suppression of high-temperature growth. (A) The interaction of Rad24 with various truncated Mks1 fragments. *rad24:GFP* cells expressing various Mks1 fragments were cultured in YES medium at 30°C and sifted to 39°C. After 2-hour incubation at 39°C, their cell lysate was subjected to immunoprecipitation using the anti-GFP antibodies (IP: GFP), and co-purified Mks1-FLAG was analyzed by immunoblotting. A green box in the schematic diagram of the Mks1 fragments indicates the region required for the Mks1-Rad24 interaction. (B) The growth of the strains expressing the various Mks1 fragments at high temperatures. The indicated *mks1* mutant strains were grown in YES medium at 30°C, and their serial dilutions were spotted onto YES agar plates for growth assay at 30°C and 39°C. (C) Amino acid sequence alignment of the C-terminal conserved region of Mks1 in *S. pombe*, *S. japonicus*, *A. nidulans,* and *N. crassa*. The sequence alignment was performed using the CLUSTALW program (https://www.genome.jp/tools-bin/clustalw). Asterisks, identical amino acids; single and double dots, weakly and strongly similar amino acids, respectively. The conserved sequence stretches are shown in a red box. (D, E) The C-terminal conserved region of Mks1 is indispensable for its function of suppressing cell growth at high temperatures. The growth of the indicated *mks1* mutant cells at 39°C was examined as in (B). A blue box in a schematic diagram of the Mks1 fragments in (D) indicates the conserved C-terminal region. (F) A model illustrating the negative regulation of cell growth under heat stress in fission yeast.

Interestingly, strains expressing the C-terminally truncated Mks1 proteins, such as Mks1(1-480) that can bind Rad24 (Fig. 6A), showed significant growth at 39°C (Fig. 6B), suggesting that the C-terminal region of Mks1 is essential in suppressing growth at high temperatures. We noticed that residues 520-539 within this region are highly conserved among the Mks1 orthologs in several fungi, including *Schizosaccharomyces japonicus*, *Aspergillus nidulans,* and *Neurospora crassa,* but not in *S. cerevisiae* (Fig. 6C). Thus, we next examined whether this conserved 20-residue sequence is required for the Mks1 function. Cells expressing the C-terminally truncated Mks1(1–510) that lacks this sequence stretch grew at 39°C, like the *mks1Δ* mutant (Fig. 6D). In contrast, the Mks1(1-550) protein, which retains the conserved stretch, was capable of suppressing cell growth at this temperature. We also constructed a strain expressing Mks1 without this sequence motif, Mks1(Δ520-539), and the mutant strain was found to exhibit a *mks1Δ*-like phenotype at 39°C (Fig. 6D). These results strongly suggest that the conserved C-terminal motif of Mks1 (Fig. 6C) is critical for the Mks1 function.

To further characterize this conserved C-terminal motif of Mks1, we constructed a series of alanine-substitution mutants, in which the two successive residues within this region were replaced with alanine at a time. While the temperature sensitivity of the *mks1-RD530.531AA* and *-QK538.539AA* mutants was comparable to that of the wild-type strain, the other *mks1* alanine mutants showed significant growth at 39°C (Fig. 6E). As expected, the Mks1-Rad24 interaction was not significantly affected by these alanine substitutions (Fig. S6), confirming that the Mks1 defects caused by these C-terminal mutations are independent of the Mks1-Rad24 interaction.

Altogether, these results strongly suggest that Mks1 has a conserved C-terminal motif essential for its function in suppressing cellular growth at high temperatures. The Mks1 function is also modulated by the 14-3-3 protein Rad24, which interacts with residues 201-280 of Mks1.

## Discussion

Increasing concerns over climate warming (51) include how higher ambient temperatures affect living organisms and their ecosystems. A recent study indicates that the rate of heat death in diverse ectotherms is extremely temperature-sensitive and doubles for every 1°C rise (52). Such acute temperature sensitivity is also observed in fission yeast cells under stressful temperatures; even a slight temperature increase from 37°C results in dramatically compromised proliferation and survival (Fig. 1). In this study, we have discovered that the thermal limit of fission yeast growth is significantly extended by rapamycin, a specific inhibitor of TORC1.

TORC1, a well-known growth-promoting factor that potentiates anabolic processes such as protein synthesis, has turned out to be a suppressor of proliferation at high temperatures, most likely through the Sck1 kinase (Figs. 1 and 2). A previous study in budding yeast found that the protein aggregation induced by arsenite attenuates cell growth and that this cytotoxic effect is suppressed by blocking the accumulation of protein aggregates with cycloheximide, an inhibitor of protein translation (53). It is therefore plausible that the downregulation of protein synthesis by TORC1 inhibition results in less heat stress-induced protein aggregation, enabling cells to proliferate at high temperatures. However, a recent study indicates that the rate of protein synthesis is not significantly affected by the triple deletion of the AGC-family kinases Psk1, Sck1, and Sck2 in fission yeast (54). Thus, the enhanced high-temperature growth by the *sck1Δ* mutation or TORC1 inhibition may be independent of protein synthesis, and the proteotoxic stress may not be a major cause of the growth arrest at temperatures marginally above 37°C.

It was unexpected that the thermal limit of fission yeast growth is dictated by an intrinsic cellular mechanism, TORC1-Sck1 signaling, rather than the level of heat injury that brings about physiological collapse. Moreover, the finding that the deletion of the *sck1*^+^ gene unleashes cell proliferation above 38°C implied that there may be additional genes inhibitory to high-temperature growth, including the one encoding the unknown substrate of the Sck1 kinase. Therefore, we screened a haploid null mutant library of *S. pombe* and identified four more genes whose null mutants exhibit growth even at 39°C (Fig. 3). Genetic analysis suggested that all those genes function independently of Sck1 to negatively regulate cell proliferation at high temperatures. The genes that we identified include an ortholog of budding yeast Mks1, which negatively regulates a set of genes involved in lysine biosynthesis and the TCA cycle (47, 55–57). However, our gene expression profiling analysis of the fission yeast *mks1Δ* mutant found no significant alteration to the mRNA levels of the lysine biosynthetic and TCA cycle genes (Table S1). Thus, the function of Mks1 in gene regulation may not be conserved between budding and fission yeasts, consistent with the limited sequence homology between their Mks1 orthologs (Fig. S2).

On the other hand, like its budding yeast counterpart, Mks1 in fission yeast physically interacts with the 14-3-3 protein Rad24, whose Glu-185-to-Val mutation was isolated during our screen for spontaneous heat-resistant mutants (Fig. 4). 14-3-3 proteins regulate various cellular processes through interaction with a diverse array of proteins (58, 59). Interestingly, Glu-185 in Rad24 is highly conserved among 14-3-3 proteins in divergent eukaryotic species (60), and this glutamate residue directly binds some of the 14-3-3 protein ligands (61, 62). Indeed, the *rad24-E185V* mutation attenuates the interaction of Rad24 with Mks1, implying the involvement of Rad24 Glu-185 in the Rad24-Mks1 complex formation.

We found that the Mks1 protein markedly accumulates in cells exposed to high temperatures (Fig. 5A) and that the *mks1Δ* mutation confers heat resistance on fission yeast cells (Fig. 3A). However, the Mks1 accumulation *per se* may not be responsible for the growth inhibition at high temperatures, as rapamycin promotes both Mks1 accumulation and cell proliferation above the normal permissive temperatures. On the other hand, the observed effect of rapamycin on the Mks1 protein level implies that TORC1 contributes to the regulation of Mks1, including its proteasome-dependent degradation (Fig. 5C). Several independent phosphoproteomic studies identified multiple phosphorylated serine/threonine residues in fission yeast Mks1 (63–67). Moreover, the function of budding yeast Mks1 is believed to be regulated by TORC1-dependent phosphorylation (47, 55). Thus, in addition to Sck1, fission yeast Mks1 may also be under the regulation by TORC1-dependent phosphorylation, possibly as a complex with Rad24, in the restriction of high-temperature growth (Fig. 6F).

Our screening of the mutant library also led to the identification of Dri1 as a negative regulator of cell proliferation at high temperatures (Fig. 3A). It was recently reported that Dri1 mediates heterochromatin assembly, which is compromised when Dri1 is tagged by the green fluorescent protein at the C-terminus (Dri1-GFP) (43). On the other hand, unlike the *dri1Δ* mutant, the temperature sensitivity of the Dri1-GFP strain constructed in our laboratory is comparable to that of the wild-type strain (Fig. S7); thus, suppression of the high-temperature growth by Dri1 appears to be independent of its function in heterochromatin assembly. Indeed, such dual functionality of Dri1 may be regulated by its binding partners. Dri1 interacts with Dpb4, a subunit of DNA polymerase epsilon, to regulate heterochromatin assembly (43), whereas we found that Rhs1 associates with Dri1 and suppresses high-temperature growth (Fig. 3C and D).

In summary, we have found that fission yeast is capable of growing at temperatures higher than previously thought and that such high-temperature growth is under negative regulation by cellular mechanisms, including TORC1-Sck1 (Fig. 6F). It is of great interest to understand why the thermal limit of fission yeast is set below the temperatures deleterious to cell physiology. Another significant question is whether similar negative controls, rather than direct heat damage to cellular components, determine the thermal limits in other unicellular organisms and cells in multicellular organisms. Curiously, most of the inhibitors of high-temperature growth that we identified in this study are conserved among eukaryotic species. We expect that further analyses of those factors in fission yeast and other eukaryotes will provide a novel insight into the determinants of the cellular thermal limit, which governs phenomena in multicellular systems under stressful heat, including climate warming.

## Materials and Methods

### Fission yeast strains and general methods

*S. pombe* strains used in this study are listed in Table S2. Growth media and basic techniques for *S. pombe* have been described previously (68, 69). For the strain constructions, the PCR-based method was applied as previously reported (70, 71).

### *S. pombe* growth assay

Fission yeast cells were grown in YES liquid medium, and the cultures were adjusted to the cell concentration equivalent to an optical density at 600 nm (OD_600_) of 1.0. Serial dilutions of the adjusted cultures were spotted onto agar solid media. Images were captured by the LAS-4000 system (Fujifilm, Japan). For the growth curve assay, cells were grown in YES liquid media at 30°C. Overnight cultures were adjusted to initial OD_600_ = 0.1 in fresh media, and cell density was measured at indicated time intervals. For heat treatment at 39°C, the overnight cultures were resuspended in pre-warmed fresh media.

### Cell viability assay

Cells were suspended in 1× PBS buffer, pH 7.4 (40 mM K_2_HPO_4_, 10 mM KH_2_PO_4_, 0.15 M NaCl). The cell suspension was then mixed with methylene blue (0.1 mg/ml stock solution) in a 1:1 ratio and incubated for 2 min at room temperature. Cell viability was examined under the microscope, and stained cells were counted as dead cells. At least 200 cells were counted for each experiment.

### Immunoblotting

Crude cell lysates were prepared using trichloroacetic acid (TCA) as described previously (72). Proteins were separated by SDS-PAGE, transferred to nitrocellulose membrane, and probed with primary antibodies as follows; anti-phospho-p70 S6K (1:5000; Cat. no. 9206, Cell Signaling Technology) for phosoho-Psk1 (Thr-415) detection, anti-Spc1 (1:10000) (72), anti-FLAG (1:5000; M2, Sigma-Aldrich), anti*-myc* (1:5000; 9E10, Covance), anti-HA (1:2000; 12CA5, Roche) and anti-GFP (1:2500; Cat. no. 04404, Nacalai Tesque). Anti-rabbit IgG (H+L) HRP-conjugated (1:10000; Cat. no. W4011, Promega), anti-mouse IgG (H+L) HRP-conjugated (1:10000; Cat. no. W4021, Promega), and anti-rat IgG (H+L) HRP-conjugated (1:10000; Cat. no. 112-035-003, Jackson Immuno Research) were used as secondary antibodies.

### Immunoprecipitation assay

Yeast cells were disrupted in lysis buffer (20 mM HEPES-NaOH [pH 7.5], 150 mM sodium glutamate, 10% glycerol, 0.25% Tween-20, 10 mM sodium fluoride, 10 mM p-nitrophenylphosphate, 10 mM sodium pyrophosphate, 10 mM β-glycerophosphate and 0.1 mM sodium orthovanadate), containing 1 mM PMSF and protease inhibitor cocktail (P8849, Sigma-Aldrich) with glass beads using Multi-beads Shocker (Yasui Kikai). The cell lysate was recovered by centrifugation for 15 min at 17,700 × g and the total protein concentrations of cell lysates were determined by Bradford assay. For interaction between FLAG- and myc-tagged proteins, the recovered cell lysates were incubated with anti-FLAG M2-affinity gel (Sigma-Aldrich) for 2 h at 4°C, followed by extensive washes with lysis buffer. Resultant samples were subjected to immunoblotting.

For interaction between FLAG-tagged Mks1 and GFP-tagged Rad24, the cell lysates were incubated with anti-FLAG antibodies (WAKO) for 1 h at 4°C, followed by incubation with Dynabeads Protein G (Invitrogen) for 2 h at 4°C.

### Whole-genome sequencing and data analysis

Cells were grown in YES liquid medium to exponential phase at 30°C and harvested by centrifugation. Genomic DNA was extracted and purified using NucleoBond AXG and Buffer Set III (MACHEREY-NAGEL) according to the manufacturer’s protocol. Whole-genome sequencing was performed by BGI JAPAN using a DNBseq platform, and the sequencing reads were mapped to the *S. pombe* reference genome using BWA (73), followed by variant calling using GATK HaplotypeCaller (74, 75). The mutations specifically found in the mutant strains were extracted and annotated using bcftools (76) and SnpEff (77), respectively.

### RNA extraction, quantitative RT-PCR (qPCR) and RNA-seq

Cells were grown to exponential phase at 30°C and shifted to 39°C for heat treatment experiment. Cells were harvested through filtration onto 0.45 µm mixed cellulose ester membrane (Advantec) and flash-frozen using liquid nitrogen. The cell pellets were resuspended in a preheated (65°C) solution composed of 25 µl of 10% SDS and 300 µl of acid phenol: chloroform, and acid-washed glass beads and bead buffer (75 mM NH_4_OAc and 10 mM EDTA, pH8) were then added. The samples were vigorously mixed using a vortex mixer for 1 min, and subsequently incubated at 65°C for 1 min. After repeating this process 3 times, the aqueous phase was recovered through centrifugation at 16000 ×*g* for 15 minutes and transferred into 250 µl of pre-warmed bead buffer. The samples were further subjected to centrifugation at 16000 ×*g*, 4°C for 15 minutes and the supernatant was mixed with prechilled bead buffers containing isopropanol. After centrifuging at 22000 ×*g*, 4°C for 30 minutes, the pellets were washed with RNase-free 70% ethanol, dried and dissolved in 30 µL of DEPC water.

For qPCR, the SuperScript II reverse transcriptase (Thermo Fisher Scientific) was used to synthesize cDNA according to the manufacturer’s protocols. qPCR was carried out using PowerUp SYBR Green Master mix (Thermo Fisher Scientific). The results were normalized against the housekeeping gene *act1^+^*.

For RNA-seq, cDNA libraries of two biological replicates for wild-type and *mks1*Δ strains were prepared following the previous paper (78), except the step of rRNA removal that was performed using RiboMinus Transcriptome Isolation Kit, yeast (ThermoFisher, K155003). Paired end sequencing was performed using Illumina HiSeq (300 cycles), and gene expression level was calculated using Salmon (version 1.1.0) (79) and the R package “DESeq2” (80). The reference genome sequence and gene annotation information of S. pombe were obtained from the PomBase database (81).

## Data Availability

The whole-genome sequence and RNA sequence data in this article have been deposited in the DDBJ Sequence Read Archive (DRA) under accession codes DRR458346-DRR458347 and DRR459935-DRR459938, respectively.

## Acknowledgments

We thank A Higuchi for technical assistance and D Watanabe for discussion. We are also grateful to T Toda and M Yukawa for sharing unpublished data and strains, and to T Matsumoto and the NBRP Japan for strains.

## Author Contributions

Y.M., K.O., and K.S., designed research; Y.M., F.M., Y.N., S.J.X., S.Y., I.Y., F.S., A. H. O., M.T., S.K., and Y.A. performed research; Y.M., F.M., Y.N., S.J.X., I.Y., F.S., A. H. O., M.T., analyzed data; and Y.M., and K.S., wrote the paper.

