## Supplementary information for "Rapamycin-sensitive mechanisms confine the growth of fission yeast below the temperatures detrimental to cell physiology"

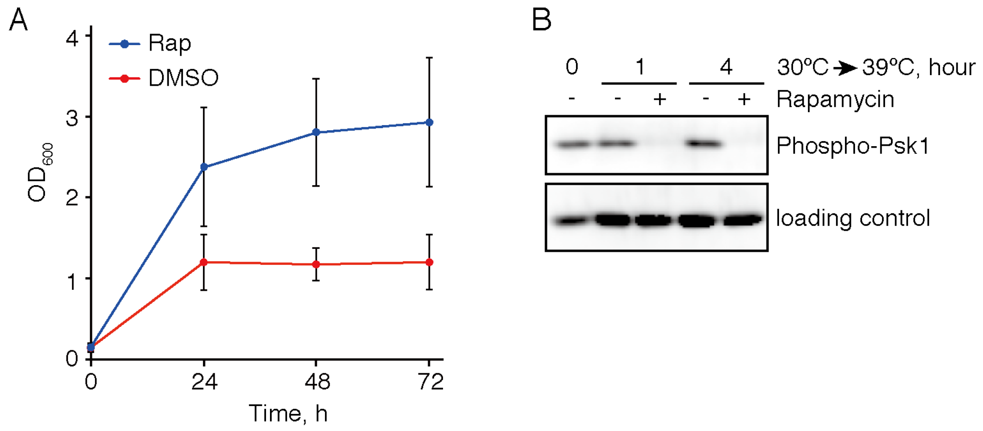


**Fig. S1. TORC1 suppression by rapamycin allows *S. pombe* cells to grow at 39˚C**

(A) Wild-type cells were grown in YES liquid medium at 30˚C. At 0 hour, the initial OD was adjusted to OD600 = 0.1 ± 0.05, and the cell growth in the presence (Rap) and absence (DMSO) of 200 ng/ml rapamycin at 39˚C was monitored by measuring OD600 at indicated time points. Data are presented as means ± s.d. from three independent experiments. (B) Wild-type cells were grown in YES medium at 30˚C and shifted to 39˚C. Cells were harvested at indicated time points and their cell lysate was subjected to immunoblotting using anti-phospho-S6K1 (Phospho-Psk1) and anti-Spc1 (loading control) antibodies.

```
S. pombe        1 -MAKRVVSPNP---MLSLETENIAKMGTLTADSLGSMWNVFTKCAENLENGRRLENISWR 56
S. cerevisiae   1 MSREAFDVPNIGTNKFLKVTPNLFTPERLNLFDDVELYLTLIKASKCVEQGERLHNISWR 60
                     : .  **     :   * *: .   *.  .  .:: .: *.:: :*:*.**.*****

S. pombe       57 LWYREAMMSDANDACQIACQESSVPDLSSSCDSINSTVESDAGHVVDSNSFNRIDNVSVN 116
S. cerevisiae  61 ILN-KAVLKEHNINRSKKRDGVKNIYYVLNPNNKQPIKPKQAAVKQPPLQKANLPPTTAK 119
                  :   :*::.: *   .   :  .      . :. :.   .:*.    . .  .:  .:.:

S. pombe      117 APIVNEVSLQAPMKGGSHSSIVRPQAKRSSSRLLSTDAFSQFISSFSPPEKPSMKDLALF 176
S. cerevisiae 120 QNVLTRPMTSPAIAQGAHDRSLDNPNSTNNDVKNDVAPNRQFSKSTTSGLFSNFADKYQK 179
                    ::..   ...:  *:*.  :    . ...   ..  .  ** .* :.   ..: *

S. pombe      177 HGNKSPSSKETIPKVSNSNSSDTSTDDQAYLNVSVSENEHADRSIKTHSSAPPLNQQKSS 236
S. cerevisiae 180 MKNVNHVANKEEPQTIITGFDTSTVITKKPLQSRRSRSPFQHIGDMNMNCIDNETSKSTS 239
                    * .  :::  *:.  :. . ::.  :  *:   *.. . . .  . ..    ..:.:*

S. pombe      237 TSVNDAVSKAIQLVKSTSDLG-SLSTNTSSTAQKNKSRKPTKSFSDAVAAASRAKELEKE 295
S. cerevisiae 240 PTLENMGSRKSSFPQKESLFGRPRSYKNDQNGQLSLSKTSSRKGKNKIFFSSEDEDSDWD 299
                  .::::  *:  .: :. * :* . * :.....* . *:..::. .: :  :*. :: : :

S. pombe      296 KLSKVTVTPDDSTSIIRGFDPRLPVLSTKNVSHEEKSHSVQDDKSKQLLKPNPTQPYFYL 355
S. cerevisiae 300 SVSNDSEFYADEDDEE--YDDYNEEEADQYYRRQWDKLLFAKNQ--QNLDSTKSSVSSAN 355
                  .:*: :    *. .    :*      : :   :: ..  . .::  * *... :.

S. pombe      356 RGSFDTGHSASSSICSAAVDSESVNSADFPHRTVYDPLPTHAIAES--ISQENVIEDDDD 413
S. cerevisiae 356 TINSNTSHDPVRKSLLSGLFLSEANSNSNNHNTAHSEYASKHVSPTPQSSHSNIGPQPQQ 415
                    . :*.*..  .   :.:  ...** .  *.*.:.  .:: :: :   *:.*:  : ::

S. pombe      414 DDDAWVSVDEAESPHFTK-RSPAPYLSKSFRNSALSLLLSQDEKLQHDVRSASSAALNLP 472
S. cerevisiae 416 NPPSANGIKQQKPSLKTSNVTALASLSPPQPSNNERLSMDIQKDFKTDNESNHLYESNAP 475
                  :  :  .:.: :..  *.  :. . ** .  ..   * :. ::.:: * .*      * *

S. pombe      473 RDTDIKATPNLSQSGNINSDNSDLSNYPYNDYRVYRMSHCSSNSNKVLASEISESLRRDL 532
S. cerevisiae 476 LTAQTILPTALSTHMFLPNNIHQQRMAIATGSNTRHRFSRRQSMDIPSKNRNTGFLKTRM 535
                    ::   .. **    : .:  :      .. .. :     .. :    .. :  *:  :

S. pombe      533 LWERRQKAAMNSAVLRRQSSQSSGANDDKEVRNRKVVEKGFANDCSVW-            580
S. cerevisiae 536 EISEEEKMVRTISRLDNTSIANSNGNGNDDTSNQRTEALGRKTSNGGRI            584
                    ...:* . . : * . *  .*..*.:.:. *::.   *  .. .
```

**Fig. S2. Amino acid sequence alignment of Mks1 in fission yeast and budding yeast**

Amino acid sequence alignment of the Mks1 protein in *S. pombe* and *S. cerevisiae*. The sequence alignment was performed using the CLUSTALW program. Asterisks, identical amino acids; single and double dots, weakly and strongly similar amino acids, respectively. The highly conserved region is shown in a red box.

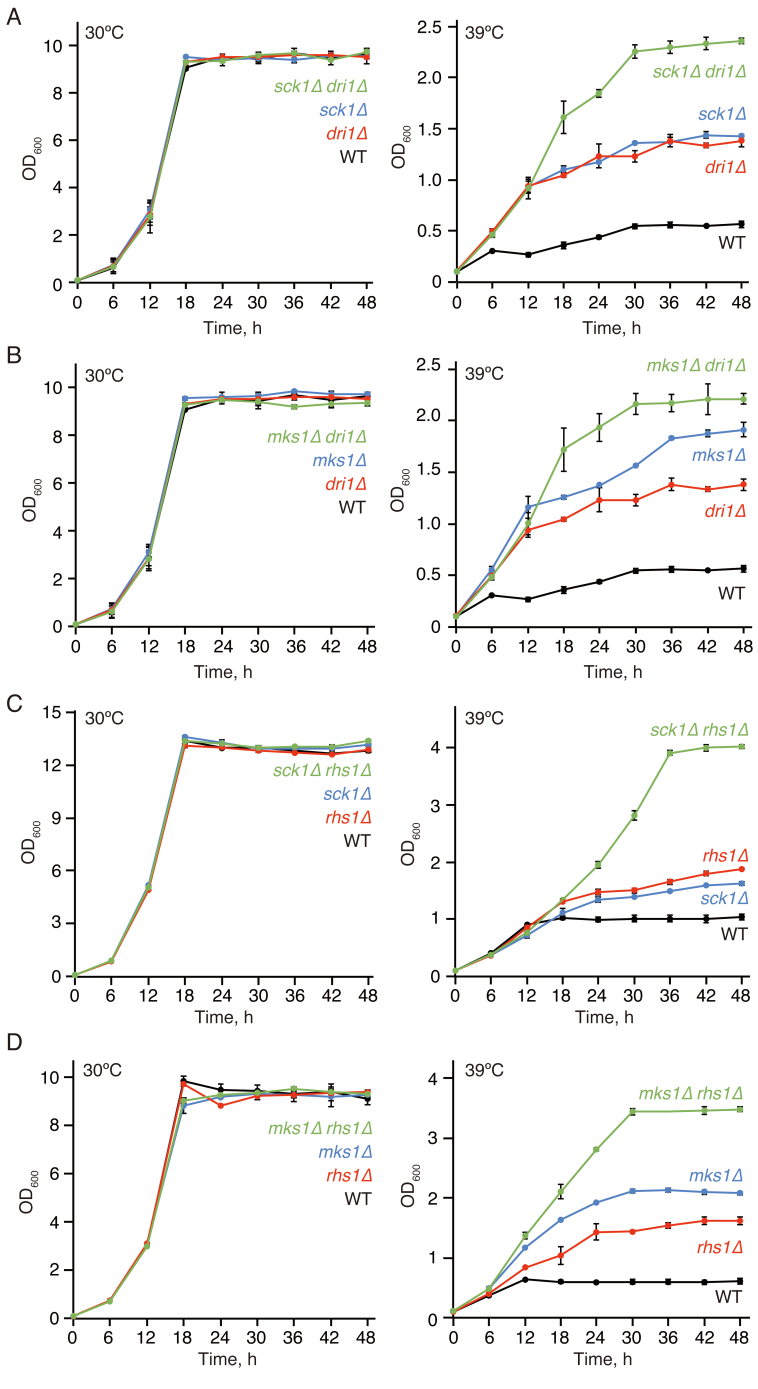


**Fig. S3.** ***nrp1*⁺ and *rhs1*⁺ act independently of *sck1*⁺ and *mks1*⁺**

(A-D) Wild-type (WT) and indicated mutant cells were in YES medium at 30˚C. The initial OD was adjusted to OD600 = 0.1 ± 0.05, and their growth at 30℃ and 39℃ was monitored by measuring OD600 every 6 hours. Data are presented as means ± s.d. from three independent experiments.

```
Rad24    1 MSTTSREDAVYLAKLAEQAERYEGMVENMKSVASTDQELTVEERNLLSVAYKNVIGARRA 56
Rad25    1 MS-NSRENSVYLAKLAEQAERYEEMVENMKKVACSNDKLSVEERNLLSVAYKNIIGARRA 59
           ** .***::************** ******.**.::::*:*************:******

Rad24   61 SWRIVSSIEQKEESKGNTAQVELIKEYRQKIEQELDTICQDILTVLEKHLIPNAASAESK 120
Rad25   60 SWRIISSIEQKEESRGNTRQAALIKEYRKKIEDELSDICHDVLSVLEKHLIPAATTGESK 119
           ****:*********:*** *. ******:***:**. **:*:*:******** *::.***

Rad24  121 VFYYKMKGDYYRYLAEFAVGEKRQHSADQSLEGYKAASEIATAELAPTHPIRLGLALNFS 180
Rad25  120 VFYYKMKGDYYRYLAEFTVGEVCKEAADSSLEAYKAASDIAVAELPPTDPMRLGLALNFS 179
           *****************:***  :.:**.***.*****:**.***.**.*:*********

Rad24  181 VFYYEILNSPDRACYLAKQAFDEAISELDSLSEESYKDSTLIMQLLRDNLTLWTSDAEYS 240
Rad25  180 VFYYEILDSPESACHLAKQVFDEAISELDSLSEESYKDSTLIMQLLRDNLTLWTSDAEYN 239
           *******:**: **:****.***************************************.

Rad24  241 AAAAGGNTEGAQENAPSNAPEGEAEPKADA-                              270
Rad25  240 QSAKEEAPAAAAASENEHPEPKESTTDTVKA                              270
            :*    . .*  .  .:.   *: ..:
```

**Fig. S4. Amino acid sequence alignment of Rad24 and Rad25**

Amino acid sequence alignment of Rad24 and Rad25. The sequence alignment was performed as in Fig. S2. The conserved glutamic acid is shown in red.

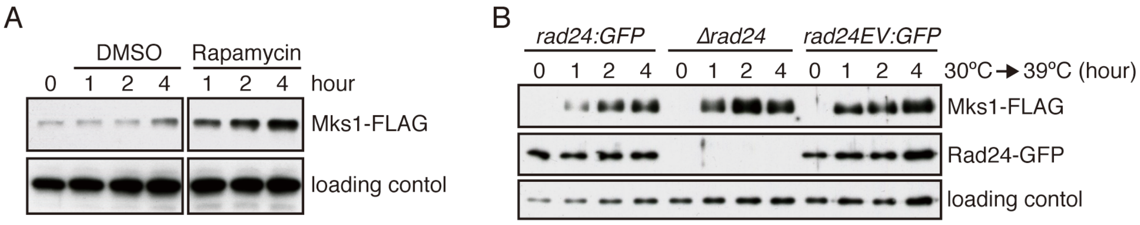


**Fig. S5. Cellular accumulation of the Mks1 protein (related to Fig. 5)**

(A) Cells were grown in YES medium at 30˚C in the presence and absence of 200 ng/ml rapamycin for indicated time, and their cell lysate was probed with anti-FLAG and anti-Spc1 (loading control) antibodies. (B) The indicated strains expressing Mks1-FLAG were cultured at 30˚C. Cells were harvested at indicated time points after shifting to 39˚C, and their cell lysate was probed with anti-FLAG, anti-GFP and anti-Spc1 (loading control) antibodies.

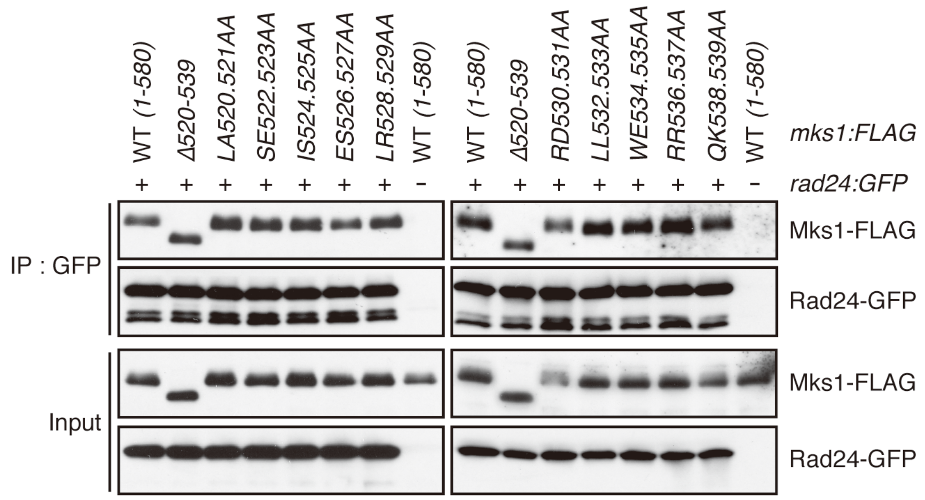


**Fig. S6. The C-terminal conserved region of Mks1 is dispensable for its interaction with Rad24**

Indicated strains were cultured in YES medium at 30˚C and sifted to 39˚C. After 2 hours of incubation at 39°C, their cell lysate was subjected to immunoprecipitation using the anti-GFP antibodies (IP: GFP), and co-purified Mks1-FLAG was analyzed by immunoblotting.

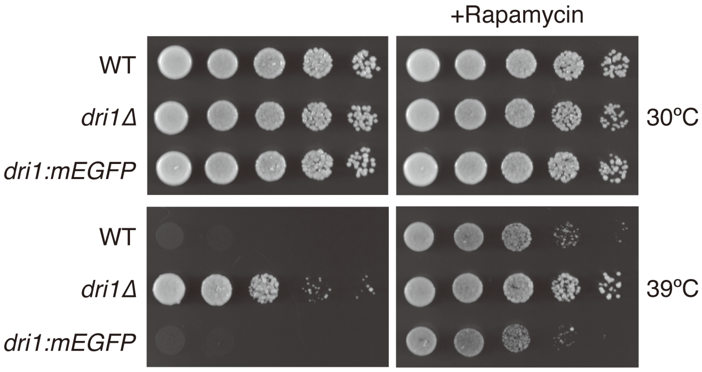


**Fig. S7. Dri1-GFP is functional in the regulation of high-temperature growth**

Indicated strains were grown in YES liquid medium at 30˚C. Their growth in the presence and absence of rapamycin (100 ng/ml) was tested at 30℃ and 39˚C by spotting serial dilutions on YES agar plates.

**Table S1. List of genes deregulated in the *mks1*Δ mutant.**

| Gene ID | Gene name | Description | $log_2$ fold change |
|---|---|---|---|
| SPCC1919.02 | pbn1 | pig-X, glycosylphosphatidylinositol-mannosyltransferase I complex subunit | 1.10 |
| SPBC21H7.04 | dbp7 | ATP-dependent RNA helicase Dbp7 | 0.85 |
| SPAC17D4.04 | trm401 | tRNA (cytosine-5-)-methyltransferase | 0.799 |
| SPAC1142.04 | noc201 | Noc complex subunit Noc201 | 0.72 |
| SPBC1734.01c | esf1 | pre-rRNA processing protein Esf1 | 0.65 |
| SPBC4F6.14 | nop4 | RNA-binding protein Nop4 | 0.64 |
| SPAC22F8.09 | rrp16 | rRNA processing protein Rrp16 | 0.57 |
| SPAC57A7.06 | utp14 | U3 snoRNP protein Utp14 | 0.54 |
| SPAC6G9.02c | nop9 | pumilio family RNA-binding protein Nop9 | 0.47 |
| SPAC23H4.09 | cdb4 | endonuclease Cdb4 | 0.44 |
| SPAC22G7.06c | ura1 | carbamoyl-phosphate synthase (glutamine hydrolyzing), aspartate carbamoyltransferase Ura1 | -0.42 |
| SPCC794.12c | mae2 | malic enzyme, malate dehydrogenase (oxaloacetate decarboxylating), Mae2 | -0.44 |
| SPAC1071.10c | pma1 | plasma membrane P-type proton exporting ATPase, P3-type Pma1 | -0.46 |
| SPCC1739.13 | ssa2 | Hsp70 family heat shock protein Ssa2 | -0.50 |
| SPAPB18E9.05c | | dubious | -0.51 |
| SPAC19G12.10c | cpy1 | vacuolar carboxypeptidase Y | -0.52 |
| SPAC513.01c | eft201 | translation elongation factor 2 (EF-2) Eft2, A | -0.52 |
| SPAPB15E9.01c | pfl2 | cell surface glycoprotein, flocculin Pfl2 | -0.54 |
| SPCC1259.01c | rps1802 | 40S ribosomal protein S18 | -0.54 |
| SPAC19A8.04 | erg5 | C-22 sterol desaturase Erg5 | -0.58 |
| SPAC821.09 | eng1 | cell septum surface endo-1,3-beta-glucanase Eng1 | -0.58 |
| SPMIT.10 | atp9 | F1-FO ATP synthase subunit 9 | -0.59 |
| SPAC1A6.04c | plb1 | phospholipase B homolog Plb1 | -0.63 |
| SPCC1840.02c | bgs4 | cell wall and secondary septum 1,6 branched 1,3-beta-glucan synthase catalytic subunit Bgs4 | -0.64 |
| SPCC18.14c | rpp0 | 60S acidic ribosomal protein Rpp0 | -0.64 |
| SPCC18.01c | adg3 | beta-glucosidase Adg3 | -0.67 |
| SPAPB1E7.12 | rps602 | 40S ribosomal protein S6 | -0.69 |
| SPAC17A5.03 | rpl301 | 60S ribosomal protein L3 | -0.70 |
| SPAC20G4.06c | adf1 | actin depolymerizing factor, cofilin | -0.71 |
| SPCC1281.06c | ole1 | acyl-coA desaturase | -0.72 |
| SPCC162.07 | ent1 | epsin | -0.73 |
| SPAPB1E7.04c | cts2 | chitinase Cts2 | -0.74 |

| | | | |
|---|---|---|---|
| SPAC26F1.07 | | NADPH-dependent aldo-keto reductase | -0.76 |
| SPBC3D6.02 | but2 | But2 family protein But2, similar to cell surface molecules | -0.81 |
| SPBC4F6.09 | str1 | plasma membrane siderophore-iron transmembrane transporter Str1 | -0.81 |
| SPBC839.16 | thf1 | C1-5,6,7,8-tetrahydrofolate (THF) synthase, trifunctional enzyme Thf1 | -0.84 |
| SPBC56F2.09c | arg5 | arginine specific carbamoyl-phosphate synthase subunit Arg5 | -0.84 |
| SPBC1703.13c | | mitochondrial carrier, inorganic phosphate/copper | -0.88 |
| SPAC1F8.03c | str3 | plasma membrane heme transmembrane transporter Str3 | -0.90 |
| SPBC1105.05 | exg1 | cell wall glucan 1,6-beta-glucosidase Exg1 | -0.95 |
| SPCC63.14 | eis1 | eisosome assembly protein Eis1 | -1.39 |
| SPAC1F7.08 | fio1 | plasma membrane iron transport multicopper oxidase Fio1 | -1.77 |

**Table S2.** ***S. pombe*** **strains list used in this study.**

| Strain ID | Genotype | Sorce, Reference |
|---|---|---|
| CA15458 | *h-* | Lab stock |
| CA17743 | *h- fkh1::natR* | FY25848 (NBRP) |
| CA15543 | *h- mip1:13myc(hph)* | Morozumi et al., 2021 |
| CA14978 | *h- mip1R358A :13myc(hph)* | Morozumi et al., 2021 |
| CA15464 | *h- mip1R504A:13myc(hph)* | Morozumi et al., 2021 |
| CA15584 | *h- mip1Y533A:13myc(hph)* | Morozumi et al., 2021 |
| CA7595 | *h- sck1::kanMX4* | Bioneer |
| CA7581 | *h- sck2::kanMX4* | Bioneer |
| CA7587 | *h- psk1::kanMX4* | Bioneer |
| CA13644 | *h- maf1::kanMX4* | Bioneer |
| CA17550 | *h- atg13::hph* | This study |
| CA8399 | *h- sck1:5FLAG(hph)* | This study |
| CA16366 | *h- sck1K331A:5FLAG(hph)* | This study |
| CA16251 | *h- mks1::kanMX4* | Bioneer |
| CA16783 | *h- dri1::kanMX4* | Bioneer |
| CA16856 | *h- rhs1::kanMX4* | Bioneer |
| CA16779 | *h- mtq2::kanMX4* | Bioneer |
| CA16254 | *h- sck1::kanMX4 mks1::kanMX4* | Bioneer |
| CA17060 | *h- sck1::kanMX4 dri1::kanMX4* | Bioneer |
| CA17058 | *h+ mks1::kanMX4 dri1::kanMX4* | Bioneer |
| CA17068 | *h- sck1::kanMX4 rhs1::kanMX4* | Bioneer |
| CA17067 | *h- mks1::kanMX4 rhs1::kanMX4* | Bioneer |
| CA17010 | *h- dri1::kanMX4 rhs1::kanMX4* | Bioneer |
| CA16965 | *h+ dri1:13myc(hph)* | This study |
| CA16967 | *h+ rhs1:5FLAG(kanMX6)* | This study |
| CA16979 | *h+ dri1:13myc(hph) rhs1:5FLAG(kanMX6)* | This study |
| CA16255 | *h- mks1:5FLAG(kanMX6)* | This study |
| CA18038 | *h+ ura4-D18 mks1:5FLAG(kanMX6) cdc25:HA(*$ura4^+$*)* | This study |
| CA18040 | *h+ ura4-D18 mks1:5FLAG(kanMX6) cdc25:HA(*$ura4^+$*) mts2-1* | This study |
| CA16640 | *h- mks1(101-580):5FLAG(kanMX6)* | This study |
| CA16642 | *h- mks1(201-580):5FLAG(kanMX6)* | This study |
| CA16644 | *h- mks1(301-580):5FLAG(kanMX6)* | This study |
| CA16680 | *h- mks1(1-480):5FLAG(kanMX6)* | This study |
| CA16681 | *h- mks1(1-380):5FLAG(kanMX6)* | This study |
| CA16670 | *h- mks1(1-280):5FLAG(kanMX6)* | This study |
| CA16772 | *h- mks1(1-510):5FLAG(kanMX6)* | This study |
| CA16768 | *h- mks1(1-550):5FLAG(kanMX6)* | This study |

| | | |
|---|---|---|
| CA16764 | *h- mks1(Δ520-539):5FLAG(kanMX6)* | This study |
| CA17266 | *h- mks1LA520.521AA:5FLAG(kanMX6)* | This study |
| CA17268 | *h- mks1SE522.523AA:5FLAG(kanMX6)* | This study |
| CA17270 | *h- mks1IS524.525AA:5FLAG(kanMX6)* | This study |
| CA17093 | *h- mks1ES526.527AA:5FLAG(kanMX6)* | This study |
| CA17105 | *h- mks1LR528.529AA:5FLAG(kanMX6)* | This study |
| CA17095 | *h- mks1RD530.531AA:5FLAG(kanMX6)* | This study |
| CA17097 | *h- mks1LL532.533AA:5FLAG(kanMX6)* | This study |
| CA17107 | *h- mks1WE534.535AA:5FLAG(kanMX6)* | This study |
| CA17137 | *h- mks1RR536.537AA:5FLAG(kanMX6)* | This study |
| CA17139 | *h- mks1QK538.539AA:5FLAG(kanMX6)* | This study |
| CA17928 | *h- rad24(kanMX6)* | This study |
| CA17931 | *h- rad24E185V(kanMX6)* | This study |
| CA17428 | *h+ rad24::kanMX6* | FY21538 (NBRP) |
| CA18014 | *h- rad25(hph)* | This study |
| CA18384 | *h- rad25E184V(hph)* | This study |
| CA16878 | *h- rad25::kanMX4* | Bioneer |
| CA18077 | *h- mks1:5FLAG(kanMX6) rad24:mEGFP(kanMX6)* | This study |
| CA17330 | *h- rad24:mEGFP(kanMX6)* | This study |
| CA18089 | *h- mks1:5FLAG(kanMX6) rad24E185V:mEGFP(kanMX6)* | This study |
| CA17328 | *h- rad24E185V:mEGFP(kanMX6)* | This study |
| CA17922 | *h- mks1(Δ520-539):5FLAG(kanMX6) rad24:mEGFP(kanMX6)* | This study |
| CA18448 | *h+ mks1LA520.521AA:5FLAG(kanMX6) rad24:mEGFP(kanMX6)* | This study |
| CA18450 | *h- mks1SE522.523AA:5FLAG(kanMX6) rad24:mEGFP(kanMX6)* | This study |
| CA18452 | *h- mks1IS524.525AA:5FLAG(kanMX6) rad24:mEGFP(kanMX6)* | This study |
| CA18454 | *h- mks1ES526.527AA:5FLAG(kanMX6) rad24:mEGFP(kanMX6)* | This study |
| CA18456 | *h- mks1LR528.529AA:5FLAG(kanMX6) rad24:mEGFP(kanMX6)* | This study |
| CA18458 | *h- mks1RD530.531AA:5FLAG(kanMX6) rad24:mEGFP(kanMX6)* | This study |
| CA18460 | *h- mks1LL532.533AA:5FLAG(kanMX6) rad24:mEGFP(kanMX6)* | This study |
| CA18462 | *h- mks1WE534.535AA:5FLAG(kanMX6) rad24:mEGFP(kanMX6)* | This study |
| CA18464 | *h- mks1RR536.537AA:5FLAG(kanMX6) rad24:mEGFP(kanMX6)* | This study |
| CA18466 | *h- mks1QK538.539AA:5FLAG(kanMX6) rad24:mEGFP(kanMX6)* | This study |
| CA18102 | *h+ mks1(101-580):5FLAG(kanMX6) rad24:mEGFP(kanMX6)* | This study |
| CA18154 | *h- mks1(201-580):5FLAG(kanMX6) rad24:mEGFP(kanMX6)* | This study |
| CA18186 | *h- mks1(301-580):5FLAG(kanMX6) rad24:mEGFP(kanMX6)* | This study |
| CA18087 | *h+ mks1(1-480):5FLAG(kanMX6) rad24:mEGFP(kanMX6)* | This study |
| CA18188 | *h- mks1(1-380):5FLAG(kanMX6) rad24:mEGFP(kanMX6)* | This study |
| CA18156 | *h- mks1(1-280):5FLAG(kanMX6) rad24:mEGFP(kanMX6)* | This study |
| CA18083 | *h- mks1:5FLAG(kanMX6) rad24::kanMX6* | This study |
| CA17022 | *h- dri1:mEGFP(kanMX6)* | This study |
